# Do auditory mismatch responses differ between acoustic features?

**DOI:** 10.1101/2020.10.13.337006

**Authors:** HyunJung An, Shing Ho Kei, Ryszard Auksztulewicz, Jan W. Schnupp

**Affiliations:** Department of Neuroscience, City University of Hong Kong, Hong Kong; Department of Neuroscience, Max Planck Institute for Empirical Aesthetics, Frankfurt, Germany

**Author notes:** Equal contribution (Ryszard Auksztulewicz, Jan W. Schnupp). **Correspondence:** Prof. Jan W.Schnupp., Dr. Ryszard Auksztulewicz.

**Keywords:** Electroencephalography (EEG), Mismatch negativity (MMN), Predictive coding, Auditory processing, Multivariate decoding

## Abstract

Mismatch negativity (MMN) is the electroencephalographic (EEG) waveform obtained by subtracting event-related potential (ERP) responses evoked by unexpected deviant stimuli from responses evoked by expected standard stimuli. While the MMN is thought to reflect an unexpected change in an ongoing, predictable stimulus, it is unknown whether MMN responses evoked by changes in different stimulus features have different magnitudes, latencies, and topographies. The present study aimed to investigate whether MMN responses differ depending on whether sudden stimulus change occur in pitch, duration, location or vowel identity respectively.

To calculate ERPs to standard and deviant stimuli, EEG signals were recorded in normal-hearing participants (N=20; 13 males, 7 females) who listened to roving oddball sequences of artificial syllables. In the roving paradigm, any given stimulus is repeated several times to form a standard, and then suddenly replaced with a deviant stimulus which differs from the standard. Here, deviants differed from preceding standards along one of four features (pitch, duration, vowel or interaural level difference). The feature levels were individually chosen to match behavioral discrimination performance.

We identified neural activity evoked by unexpected violations along all four acoustic dimensions. Evoked responses to deviant stimuli increased in amplitude relative to the responses to standard stimuli. A univariate (channel-by-channel) analysis yielded no significant differences between MMN responses following violations of different features. However, in a multivariate analysis (pooling information from multiple EEG channels), acoustic features could be decoded from the topography of mismatch responses, although at later latencies than those typical for MMN. These results support the notion that deviant feature detection may be subserved by a different process than general mismatch detection.

## 1. Introduction

Neural activity is typically suppressed in response to expected stimuli and enhanced following novel stimuli (Carbajal and Malmierca, 2018). This effect is often summarized as a mismatch response, calculated by subtracting the neural response waveform to unexpected deviant stimuli from the response to expected standard stimuli. The mismatch negativity (MMN) is such a difference waveform based on event-related potentials (ERPs). In humans, the MMN typically peaks at latencies between 150 ms and 250 ms following stimulus onset (Garrido et al., 2008). The principal neural sources of the MMN are thought to be superior temporal regions adjacent to the primary auditory cortex, as well as frontoparietal areas (Doeller et al., 2003; Chennu et al., 2013). Initially, the MMN was interpreted as a correlate of pre-attentive encoding of physical features between standard and deviant sounds (Doeller et al., 2003). However, more recent studies have led to substantial revisions of this hypothesis, and currently, the most widely accepted explanation of the MMN is that it reflects a prediction error response.

An important theoretical question remains whether mismatch signaling has a domain-general or domain-specific (feature-dependent) implementation in the auditory processing pathway. A recent study using invasive recordings from the cortical surface (Auksztulewicz et al., 2018) demonstrated that neural mechanisms of predictions regarding stimulus contents (“what”) and timing (“when”) can be dissociated in terms of their topographies and latencies throughout the frontotemporal network, and that activity in auditory regions is sensitive to interactions between different kinds of predictions. Additionally, biophysical modeling of the measured signals has shown that predictions of contents and timing are best explained either by short-term plasticity or by classical neuromodulation respectively, suggesting separable mechanisms for signaling different kinds of predictions. However, these dissociations might be specific to predictions of contents vs. timing, which may have fundamentally different roles in processing stimulus sequences (Friston and Buzsaki, 2016).

Interestingly, an earlier magnetoencephalography (MEG) study by Phillips et al. (Phillips et al., 2015) provided evidence for a hierarchical model, whereby violations of sensory predictions regarding different stimulus contents were associated with similar response magnitudes in auditory cortex, but different connectivity patterns at hierarchically higher levels of the frontotemporal network. This result is consistent with the classical predictive coding hypothesis in which reciprocal feedforward and feedback connections at the lower levels of the hierarchy are thought to signal prediction errors and predictions regarding simple sensory features, but hierarchically higher levels are thought to signal more complex predictions and prediction errors, integrating over multiple features (Kiebel et al., 2008). However, in the Phillips et al. study (Phillips et al., 2015), physical differences between deviants and standards were not behaviorally matched across different features or participants, raising the possibility that differences in frontal activity might to some extent be explained by differences in stimulus salience (Shiramatsu and Takahashi, 2018). Furthermore, a recent study investigating the MMN to acoustic violations along multiple independent features in the rat auditory cortex (An et al., 2020) revealed that the topography of MMN signals was highly diverse across not only acoustic features, but also individual rats, suggesting that the spatial resolution of non-invasive methods such as EEG or MEG might not be sufficient for mapping more subtle differences between mismatch responses to violations of different features.

The present study employed a roving oddball paradigm in which auditory deviants could differ from preceding standards along one of four independent acoustic features. Unlike in the previous study of the MMN to multiple acoustic features (Phillips et al., 2015) or in previous roving paradigms (Garrido et al., 2008), we systematically matched stimulus differences for all features and participants based on individual behavioral performance. Our primary goal was to test whether mismatch responses to violations of different features differ in magnitude or latency. In addition to testing the effects of acoustic feature on the MMN time-course in a mass-univariate analysis (i.e., on an electrode-by-electrode basis), we also aimed at decoding acoustic features from differences in MMN topography in a multivariate analysis (i.e., pooling signals from multiple electrodes).

## 2. Materials and Methods

### 2.1 Participants

Twenty volunteers (13 males and 7 females; mean age 23.9 years old) enrolled in the study upon written informed consent. All participants self-reported as having normal hearing and no history of neurological disorders, and all but two were right-handed. All experimental procedures were approved by the Human Subjects Ethics Sub-Committee of the City University of Hong Kong.

### 2.2 Stimuli

In this experiment, we manipulated two consonant-vowel (CV) syllable stimuli, /ta/ and /ti/ (Retsa et al., 2018), along four independent acoustic features: duration, pitch, interaural level difference (ILD) or vowel (An et al., 2020). Prior to the EEG recording, per participant, we estimated the feature interval yielding ~80% behavioral performance by employing a 1-up-3-down staircase procedure. In each staircase trial, two out of three stimuli, chosen at random, were presented at a mean level of a given feature (e.g., a 50/50 vowel mixture or a 0 dB ILD) while the third stimulus was higher or lower than the mean level by a certain interval. Participants had to indicate which stimulus was the “odd one out”. Following three consecutive hits, the interval decreased by 15%; following a mistake, the interval increased by 15%. Each participant performed 30 staircase trials for each feature. For the roving oddball stimulus sequences, the stimulus duration was set to 120 ms and the inter-stimulus intervals (ISIs) were fixed at 500 ms. Stimuli formed a roving oddball sequence: after 4-35 repetitions of a given stimulus (forming a standard), it was replaced with another (deviant) stimulus, randomly drawn from the set of 5 possible levels (Figure 1. (A)). Roving oddball sequences corresponding to different features were administered in separate blocks, in a randomized order across participants. The total number of stimuli in each block was approximately 2000, including 200 deviant stimuli and 200 corresponding (immediately preceding) standards.

**Figure 1.**
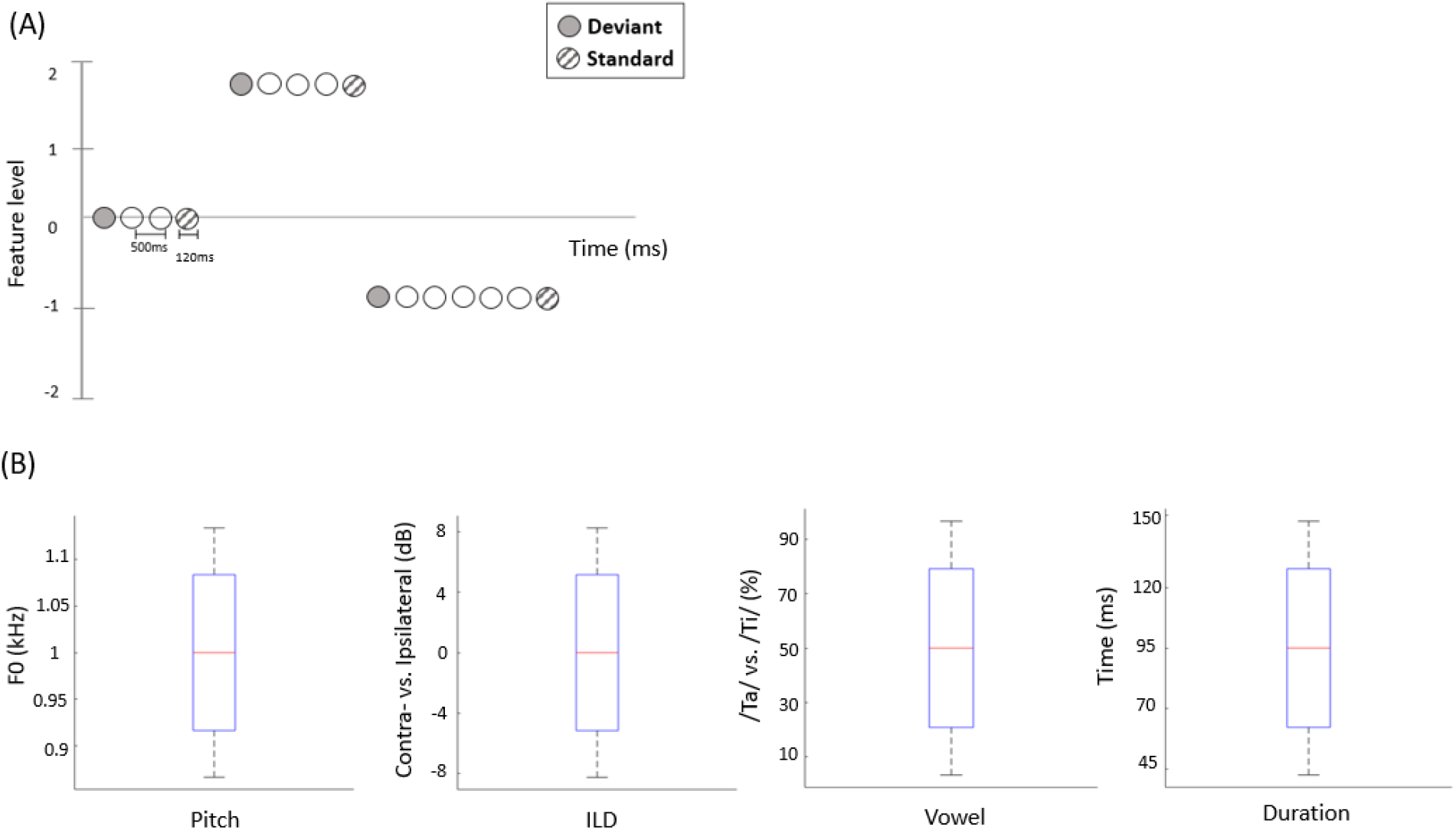
(A) Schematic representation of the stimulation sequences. The first stimulus in each train (solid circles) represents a deviant sound, while the last (hatched circles) represents a standard sound. (B) The range of each acoustic feature used to construct stimuli in the EEG experiment. Red line indicates median value of each feature (across participants), blue bars and black whiskers represent mean and SD of upper and lower ranges across participants.

### 2.3 Experimental procedure

We recorded signals from 64 EEG channels in a 10-20 system using an ANT Neuro EEG Sports amplifier. EEG channels were grounded at the nasion and referenced to the Cpz electrode. Participants were seated in a quiet room and fitted with Brainwavz B100 earphones, which delivered the audio stimuli via a MOTU Ultralite MK3 USB soundcard at 44.1 kHz. EEG signals were pre-processed using the SPM12 Toolbox for MATLAB. The continuous signals were first notch-filtered between 48 and 52 Hz and band-pass filtered between 0.1 and 90 Hz (both filters: 5th order zero-phase Butterworth), and then downsampled to 300 Hz. Eye blinks were automatically detected using the Fp1 channel, and the corresponding artefacts were removed by subtracting the two principal spatiotemporal components associated with each eye blink from all EEG channels (Ille et al., 2002). Then, data were re-referenced to the average of all channels, segmented into epochs ranging from - 100 ms before to 400 ms after each stimulus onset, baseline-corrected to the average pre-stimulus voltage, and averaged across trials to obtain ERPs for deviants and standards for each of the four acoustic features.

### 2.4 Data analyses

First, to establish the presence of the MMN response, we converted the EEG time-series into 3D images (2D spatial topography × 1D time-course) and entered them into a general linear model (GLM) with two factors (random effect of mismatch: deviant vs. standard; fixed effect of participant), corresponding to a paired t-test. Statistical parametric maps were thresholded at an uncorrected p < 0.005, and the resulting spatiotemporal clusters of main effects were tested for statistical significance at the family-wise error corrected threshold pFWE < 0.05, taking into account the spatiotemporal correlations and multiple comparisons across channels and time points. This analysis confirmed that EEG amplitudes differed significantly between deviants and standards when pooling over all the acoustic features tested (Figure 2. (A)). Specifically, the central EEG channels showed a significant mismatch response between 115 and 182 ms (cluster-level pFWE < 0.001, Tmax = 3.94), while posterior channels showed a significant mismatch response between 274 and 389 ms (cluster-level pFWE < 0.001, Tmax = 5.46), within the range of a P3b component.

**Figure 2.**
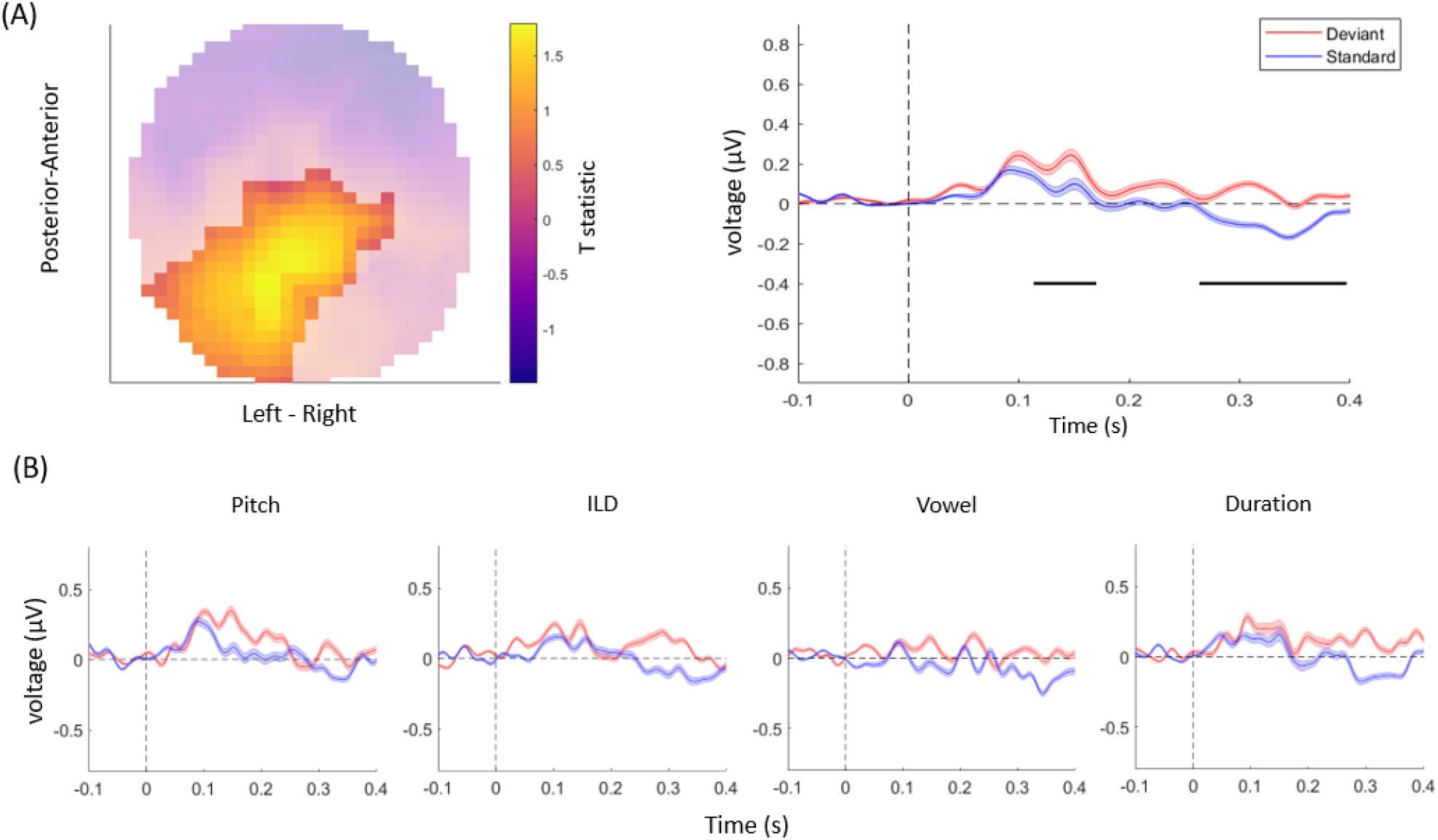
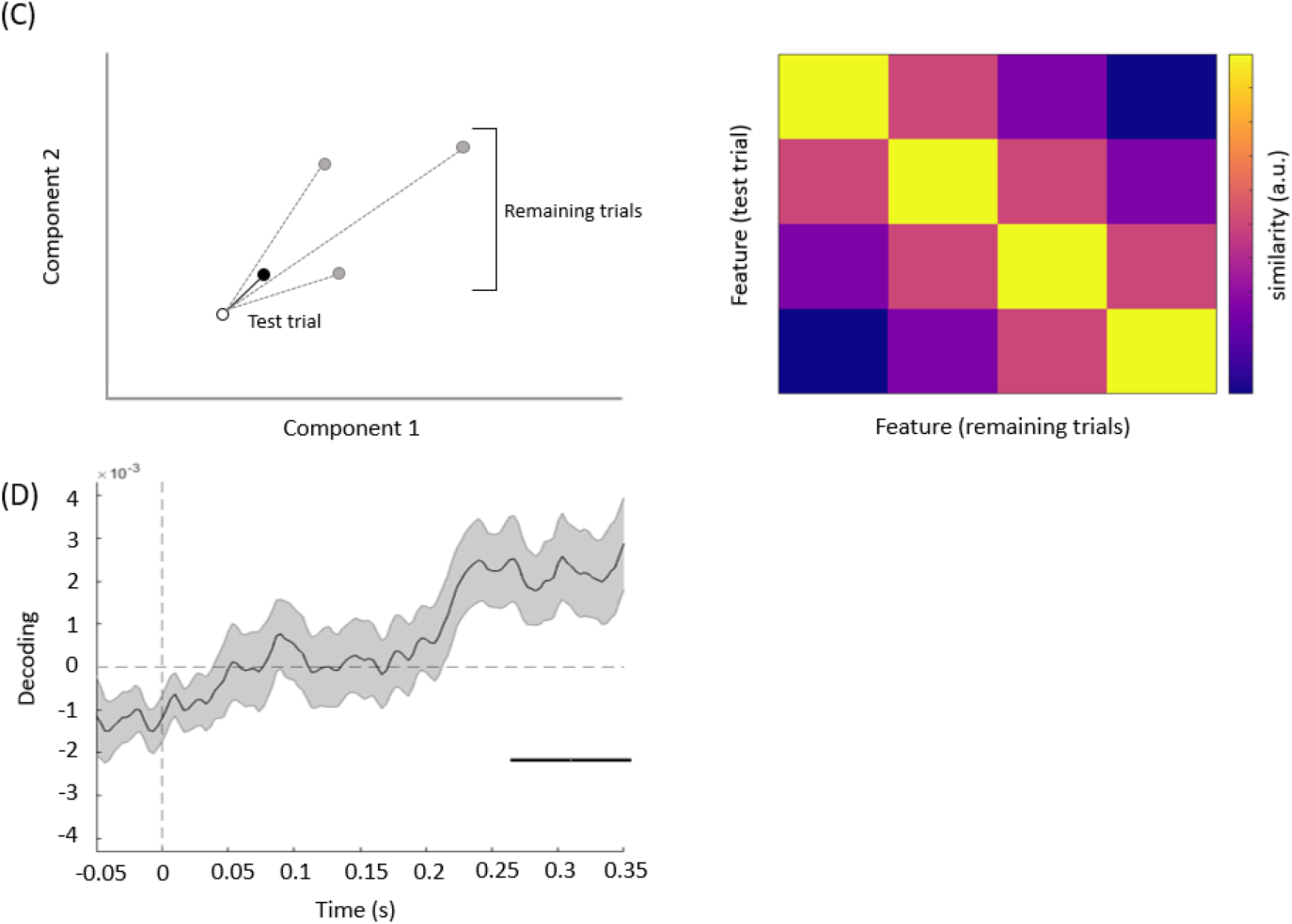
(A) The topography (left) and time-course (right) of the mismatch response. The highlighted topography cluster represents the significant difference between deviants and standards. Based on this cluster, the average waveform of the evoked response is plotted separately for auditory standards (blue) and deviants (red). The horizontal bars (black) indicate time points with a significant difference between deviants and standards. Shaded areas denote SEM across participants. (B) The average response to acoustic standards (blue) and deviants (red) for different feature conditions, extracted from the same cluster as in (A). No interaction effects were significant after correcting for multiple comparisons across channels and time points. (C) Decoding methods. Left panel: for each trial, we calculated the Mahalanobis distance, based on multiple EEG components (here shown schematically for two components), between the mismatch response in a given (test) trial (empty circle) and the average mismatch responses based on the remaining trials (black circle: same feature as test trial; grey circles: different features). Right panel: after averaging the distance values across all trials, we obtained 4 by 4 similarity matrices between all features, such that high average Mahalanobis distance corresponded to low similarity between features. Based on these matrices, we summarized feature decoding as the difference between the diagonal and off-diagonal terms. (D) Multivariate analysis. The average time course of the decoding of acoustic features based on single-trial mismatch response. The gray-shaded area denotes the SEM across participants, and the horizontal bar (black) shows the significant time window.

Then, to test whether MMN amplitudes differed between stimulus features, ERP data were entered into a flexible-factorial GLM with one random factor (participant) and two fixed factors (mismatch: deviant vs. standard; feature: pitch, duration, ILD, and vowel), corresponding to a repeated-measures 2 × 4 ANOVA. Statistical significance thresholds were set as above. The results showed no significant interaction effects between mismatch and acoustic features (all cluster-level pFWE > 0.05) (Figure 2. (B)).

Finally, to test whether mismatch responses can be used to decode the violated acoustic features, we subjected the data to a multivariate analysis. Prior to decoding, we calculated single-trial mismatch response signals by subtracting the EEG signal evoked by each standard from the signal evoked by the subsequent deviant. Data dimensionality was reduced using PCA (principal component analysis), resulting in spatial principal components (describing channel topographies) and temporal principal components (describing voltage time-series), sorted by the ratio of explained variance. Only those top components which, taken together, explained 95 % of the original variance, were retained for further analysis. In decoding acoustic features, we adopted a sliding window approach, integrating over the relative voltage changes within a 100 ms window around each time-point (Wolff et al., 2020). To this end, per channel and trial, the time segments within 100 ms of each analysed time-point were down-sampled by binning the data over 10 ms bins, resulting in a vector of 10 average voltage values per component. Next, the data were de-meaned by removing the component-specific average voltage over the entire 100 ms time window from each component and time bin. These steps ensured that the multivariate analysis approach was optimised for decoding transient activation patterns (voltage fluctuations around a zero mean) at the expense of more stationary neural processes (overall differences in mean voltage) (Wolff et al., 2020).

The binned single-trial mismatch fluctuations were then concatenated across components for subsequent leave-one-out cross-validation decoding. Per trial and time point, we calculated the Mahalanobis distance (De Maesschalck et al., 2000) (scaled by the noise covariance matrix of all components) between the vector of concatenated component fluctuations of this trial (test trial) and four other vectors, obtained from the remaining trials, and corresponding to the concatenated component fluctuations averaged across trials, separately for each of the four features. The resulting Mahalanobis distance values were averaged across trials, separately for each acoustic feature, resulting in 4 × 4 distance matrices. These distance matrices were summarized per time point and participant as a single decoding estimate, by subtracting the mean off-diagonal from diagonal terms (Figure. 2(C)).

## 3. Results

Taken together, in this study we tested whether auditory mismatch responses are modulated by violations of independent acoustic features. First, consistent with previous literature (Doeller et al., 2003; Garrido et al., 2008), we observed overall differences between the ERPs evoked by deviant stimuli vs. standard stimuli, in a range typical for MMN responses as well as at longer latencies (Figure 2. (A)). Although the ERP time-courses differed between deviant and standard stimuli when pooling over violations of different acoustic features, a univariate (channel-by-channel) analysis revealed no significant differences in the amplitudes or time-courses of mismatch responses between independent stimulus features (Figure 2. (B)). These results are consistent with a previous study (Phillips et al., 2015) which found that multiple deviant stimulus features (frequency, intensity, location, duration, and silent gap) were not associated with differences in activity in the auditory regions, but instead were reflected in more distributed activity patterns (frontotemporal connectivity estimates).

The resulting decoding time-courses of each participant were entered into a GLM and subject to one-sample t-tests, thresholded at an uncorrected p < 0.05 and correcting for multiple comparisons across time points at a cluster-level pFWE < 0.05. In this analysis, significant acoustic feature decoding was observed between 247 and 350 ms relative to tone onset (cluster-level pFWE = 0.000, Tmax = 2.77). (Figure 2. (D)). Thus, when pooling information from multiple EEG channels, acoustic features could be decoded from the topography of mismatch responses, although at later latencies than typical for MMN.

## 4. Discussion

In this study, since a univariate analysis of interactions between mismatch signals and acoustic features might not be sensitive enough to reveal subtle and distributed amplitude differences between conditions, we adopted a multivariate analysis aiming at decoding the violated acoustic feature from single-trial mismatch response topographies. This demonstrated that acoustic features could be decoded from the topography of mismatch responses, although at later latencies than typical for MMN (Figure 2. (D)). A recent oddball study (Leung et al., 2012) examined ERP differences to violations of four features (frequency, duration, intensity, and interaural difference). The study found that frequency deviants were associated with a significant amplitude change in the middle latency range. This result indicated that deviant feature detection may be subserved by a different process than general mismatch detection. Although we found feature-specificity in the late latency range, these discrepancies might be explained by different stimulus types. While the previous study used simple acoustic stimuli, here we used complex syllable stimuli, possibly tapping into the later latencies of language-related mismatch responses, as compared to MMN following violations of non-speech sounds.

Our results can be explained in terms of a hierarchical deviance detection system based on predictive coding (Kiebel et al., 2008). On this account, neural responses supporting the lower and higher hierarchical stages communicate continuously through reciprocal pathways. When exposed to repetitive stimuli, the bottom-up (ascending) sensory inputs can be “explained away” by top-down (descending) connections mediating prediction signaling, resulting in weaker prediction error signaling back to the hierarchically higher regions. Substituting the predicted standard with an unpredicted deviant results in a failure of top-down suppression by prior predictions. This leads to an increased prediction error signaling back to higher regions, providing an update for subsequent predictions. In paradigms where multiple acoustic features vary independently (such as here), low-level prediction error signals need to be integrated by hierarchically higher regions across multiple neuronal populations and time points, to infer which descending prediction needs to be updated. As a result, differences between prediction errors to violations of different stimulus features might be detected later and in more distributed activity patterns than typical mismatch responses.

In conclusion, the present study identified functional dissociations between deviance detection and deviance feature detection. First, while mismatch responses were observed at latencies typical for the MMN as well as at longer latencies, channel-by-channel analyses revealed no robust differences between mismatch responses following violations of different acoustic features. However, we demonstrate that acoustic features could be decoded at longer latencies, based on fine-grained spatiotemporal patterns of mismatch responses. This finding suggests that deviance feature detection might be mediated by later and more distributed neural responses than deviance detection itself.

## 5. Author Contributions

**HyunJung An**: Formal analysis, Writing original draft, Conceptualization, Conducted experiment. **Shing Ho Kei**: Conducted experiment, Formal analysis. **Ryszard Auksztulewicz**: Formal analysis, Supervision, Project administration, Conceptualization. **Jan W.H. Schnupp**: Project administration, Conceptualization, Investigation, Supervision.

## References

An, H., Auksztulewicz, R., Kang, H., and Schnupp, J.W.H. (2020). Cortical mapping of mismatch responses to independent acoustic features. Hear Res, 107894. doi: 10.1016/j.heares.2020.107894.

Auksztulewicz, R., Schwiedrzik, C.M., Thesen, T., Doyle, W., Devinsky, O., Nobre, A.C., et al. (2018). Not All Predictions Are Equal: “What” and “When” Predictions Modulate Activity in Auditory Cortex through Different Mechanisms. J Neurosci 38(40), 8680–8693. doi: 10.1523/JNEUROSCI.0369-18.2018.

Carbajal, G.V., and Malmierca, M.S. (2018). The Neuronal Basis of Predictive Coding Along the Auditory Pathway: From the Subcortical Roots to Cortical Deviance Detection. Trends Hear 22, 2331216518784822. doi: 10.1177/2331216518784822.

Chennu, S., Noreika, V., Gueorguiev, D., Blenkmann, A., Kochen, S., Ibanez, A., et al. (2013). Expectation and attention in hierarchical auditory prediction. J Neurosci 33(27), 11194–11205. doi: 10.1523/JNEUROSCI.0114-13.2013.

De Maesschalck, R., Jouan-Rimbaud, D., and Massart, D.L. (2000). The Mahalanobis distance. Chemometrics and Intelligent Laboratory Systems 50(1), 1–18. doi: Doi 10.1016/S0169-7439(99)00047-7.

Doeller, C.F., Opitz, B., Mecklinger, A., Krick, C., Reith, W., and Schroger, E. (2003). Prefrontal cortex involvement in preattentive auditory deviance detection: neuroimaging and electrophysiological evidence. Neuroimage 20(2), 1270–1282. doi: 10.1016/S1053-8119(03)00389-6.

Friston, K., and Buzsaki, G. (2016). The Functional Anatomy of Time: What and When in the Brain. Trends Cogn Sci 20(7), 500–511. doi: 10.1016/j.tics.2016.05.001.

Garrido, M.I., Friston, K.J., Kiebel, S.J., Stephan, K.E., Baldeweg, T., and Kilner, J.M. (2008). The functional anatomy of the MMN: a DCM study of the roving paradigm. Neuroimage 42(2), 936–944. doi: 10.1016/j.neuroimage.2008.05.018.

Ille, N., Berg, P., and Scherg, M. (2002). Artifact correction of the ongoing EEG using spatial filters based on artifact and brain signal topographies. Journal of Clinical Neurophysiology 19(2), 113–124. doi: Doi 10.1097/00004691-200203000-00002.

Kiebel, S.J., Daunizeau, J., and Friston, K.J. (2008). A hierarchy of time-scales and the brain. PLoS Comput Biol 4(11), e1000209. doi: 10.1371/journal.pcbi.1000209.

Leung, S., Cornella, M., Grimm, S., and Escera, C. (2012). Is fast auditory change detection feature specific? An electrophysiological study in humans. Psychophysiology 49(7), 933–942. doi: 10.1111/j.1469-8986.2012.01375.x.

Phillips, H.N., Blenkmann, A., Hughes, L.E., Bekinschtein, T.A., and Rowe, J.B. (2015). Hierarchical Organization of Frontotemporal Networks for the Prediction of Stimuli across Multiple Dimensions. J Neurosci 35(25), 9255–9264. doi: 10.1523/JNEUROSCI.5095-14.2015.

Retsa, C., Matusz, P.J., Schnupp, J.W.H., and Murray, M.M. (2018). What’s what in auditory cortices? Neuroimage 176, 29–40. doi: 10.1016/j.neuroimage.2018.04.028.

Shiramatsu, T.I., and Takahashi, H. (2018). Mismatch Negativity in Rat Auditory Cortex Represents the Empirical Salience of Sounds. Front Neurosci 12, 924. doi: 10.3389/fnins.2018.00924.

Wolff, M.J., Kandemir, G., Stokes, M.G., and Akyurek, E.G. (2020). Unimodal and Bimodal Access to Sensory Working Memories by Auditory and Visual Impulses. J Neurosci 40(3), 671–681. doi: 10.1523/JNEUROSCI.1194-19.2019.

